# JIP3 links lysosome transport to regulation of multiple components of the axonal cytoskeleton

**DOI:** 10.1101/2020.06.24.169219

**Authors:** N.M. Rafiq, L.L. Lyons, S. Gowrishankar, P. De Camilli, S.M. Ferguson

## Abstract

Lysosome axonal transport is important for the clearance of cargoes sequestered by the endocytic and autophagic pathways. Building on observations that mutations in the JIP3 (*MAPK8IP3*) gene result in lysosome-filled axonal swellings, we analyzed the impact of JIP3 depletion on the cytoskeleton of human neurons. Dynamic focal lysosome accumulations were accompanied by disruption of the axonal periodic scaffold (spectrin, F-actin and myosin II) throughout each affected axon. Additionally, axonal microtubule organization was locally disrupted at each lysosome-filled swelling. This local axonal microtubule disorganization was accompanied by accumulations of both F-actin and myosin II. These results indicate that transport of axonal lysosomes is functionally interconnected with mechanisms that control the organization and maintenance of the axonal cytoskeleton. They have potential relevance to human neurological disease arising from JIP3 mutations as well as for neurodegenerative diseases associated with the focal accumulations of lysosomes within axonal swellings such as Alzheimer’s disease.

## Introduction

Neurons face major demands arising from their extreme size, polarity and longevity. Axons in particular stand out due to their length which requires both long-range transport for delivery of cargoes to and from distant locations combined with mechanisms to ensure structural integrity (Hammarlund et al., 2007; Lorenzo et al., 2019; Maday et al., 2014). These challenges create unique vulnerabilities that are reflected in the numerous neurodevelopmental and neurodegenerative diseases arising from defects in axonal transport and maintenance (Coleman and Hoke, 2020; Millecamps and Julien, 2013; Sleigh et al., 2019). The unique morphology and functions of axons requires specialized organization of multiple cytoskeletal components. Axonal microtubules, which provide the tracks on which motors can transport organelles and other cargoes over long distances, are polarized with their plus ends towards the distal axon and regulated by the binding of various microtubule binding proteins (Barnes and Polleux, 2009; Dent and Gertler, 2003; Kapitein and Hoogenraad, 2011; Stiess and Bradke, 2011). The membrane associated periodic actin-spectrin lattice provides structural support to ensure axon integrity and non-muscle myosin II-dependent contractility coordinates the passage of organelles through the narrow confines of axons (Costa et al., 2020; Krieg et al., 2014; Leterrier, 2021; Vassilopoulos et al., 2019; Wang et al., 2019).

The vast majority of axonal proteins are synthesized in the neuronal cell body and proximal dendritic regions, and are subsequently transported into axons to meet their structural, signaling and metabolic demands and to support synaptic transmission (McEwen and Grafstein, 1968). Conversely, efficient retrograde transport from the axon periphery back to the cell body is required for the clearance of old or damaged proteins, as well as of material taken up by endocytosis, via endocytic and autophagic pathways (Ferguson, 2018; Kulkarni et al., 2018). This transport is primarily mediated by immature lysosomes and auto-lysosomes, which have a low content of lysosomal hydrolases and whose fate is to mature into fully degradative lysosomes in cell bodies by fusing with hydrolases-enriched vesicles delivered from the trans-Golgi network (Ferguson, 2018). The massive accumulation of these organelles at the distal side of focal blocks of axonal transport reveals that such lysosomes are the major retrograde axonal cargo (Tsukita and Ishikawa, 1980).

A similar build-up of immature lysosomes is observed in axon swellings surrounding amyloid Aβ deposits at Alzheimer’s disease amyloid plaques (both in human patients and in mouse models of the disease), which are putative sites of APP processing (Blazquez-Llorca et al., 2017; Gowrishankar et al., 2015; Nixon, 2005). A link between accumulation of axonal lysosomes due to their impaired transport and amyloidogenic APP processing was further supported by studies of neurons and mice with loss of JIP3 function (Gowrishankar et al., 2017). JIP3 is a motor interacting protein which is preferentially expressed in neurons and is thought to couple cargos such as lysosomes to dynein, the microtubule minus-end directed motor (Vilela et al., 2019). JIP3 loss-of-function mutations in multiple animal species result in the build-up of lysosomes within axons (Drerup and Nechiporuk, 2013; Edwards et al., 2013; Gowrishankar et al., 2017). For example, primary cultures of mouse JIP3 knockout neurons exhibit a striking increase in the overall abundance of axonal lysosomes with focal accumulations within axonal swellings that are strikingly similar to the lysosome-filled axonal swellings observed at amyloid plaques (Gowrishankar et al., 2017). These changes raised questions about the relationship between lysosomes and the axonal cytoskeleton with potential implications for Alzheimer’s disease.

To address these questions, we used human JIP3 KO iPSC-derived cortical glutamatergic neurons that we recently established as a cellular model for investigating the impact of JIP3 depletion on neuronal cell biology (Gowrishankar et al., 2021). Surprisingly, we found that axons with lysosome-filled swellings had a massive disruption in their actin-spectrin and myosin-II lattice organization that was not restricted to just the local site of the swelling but which occurred throughout each affected JIP3 KO axon. The additional KO of JIP4 further enhanced this phenotype, consistent with an overlapping role of the two proteins (Gowrishankar et al., 2021). Axonal swellings filled with lysosomes were not static but formed and resolved over the course of several hours. Intriguingly, their formation coincided with local microtubule disorganization. Our observations support a model wherein perturbed axonal lysosome transport induced by loss of JIP3 (or JIP3 and JIP4) is closely linked to a broad disruption of the neuronal cytoskeleton.

## Results

### Global disruption of the membrane periodic skeleton in axons with lysosome-filled axonal swellings

The axonal plasma membrane is supported by an organized cytoskeletal network containing actin filaments and spectrin tetramers (Han et al., 2017; He et al., 2016; Vassilopoulos et al., 2019).This actin-spectrin network, which has been linked to axonal mechanical stability (Hammarlund et al., 2007; Krieg et al., 2014) and signaling (Zhou et al., 2019), was also recently shown to undergo transient local expansion to facilitate the transport of large cargoes in narrow axons (Wang et al., 2020). The important role of this cytoskeletal scaffold in controlling axon diameter suggested that lysosome-filled axonal swellings of JIP3 KO neurons might require major rearrangement to this network. To address this question, human induced pluripotent stem cells (iPSCs) which can be differentiated into layer 2/3 cortical glutamatergic neurons (i^3^Neurons)(Fernandopulle et al., 2018; Gowrishankar et al., 2021) were used in this study. The KO of JIP3 in this human i^3^Neuron model system robustly develops lysosome-filled axonal swellings similar to those observed in primary cultures of mouse JIP3 KO cortical neurons while overcoming practical challenges arising from the neonatal lethality in the JIP3 KO mouse model (Figure 1A-C)(Gowrishankar et al., 2021; Gowrishankar et al., 2017).

**Figure 1:**
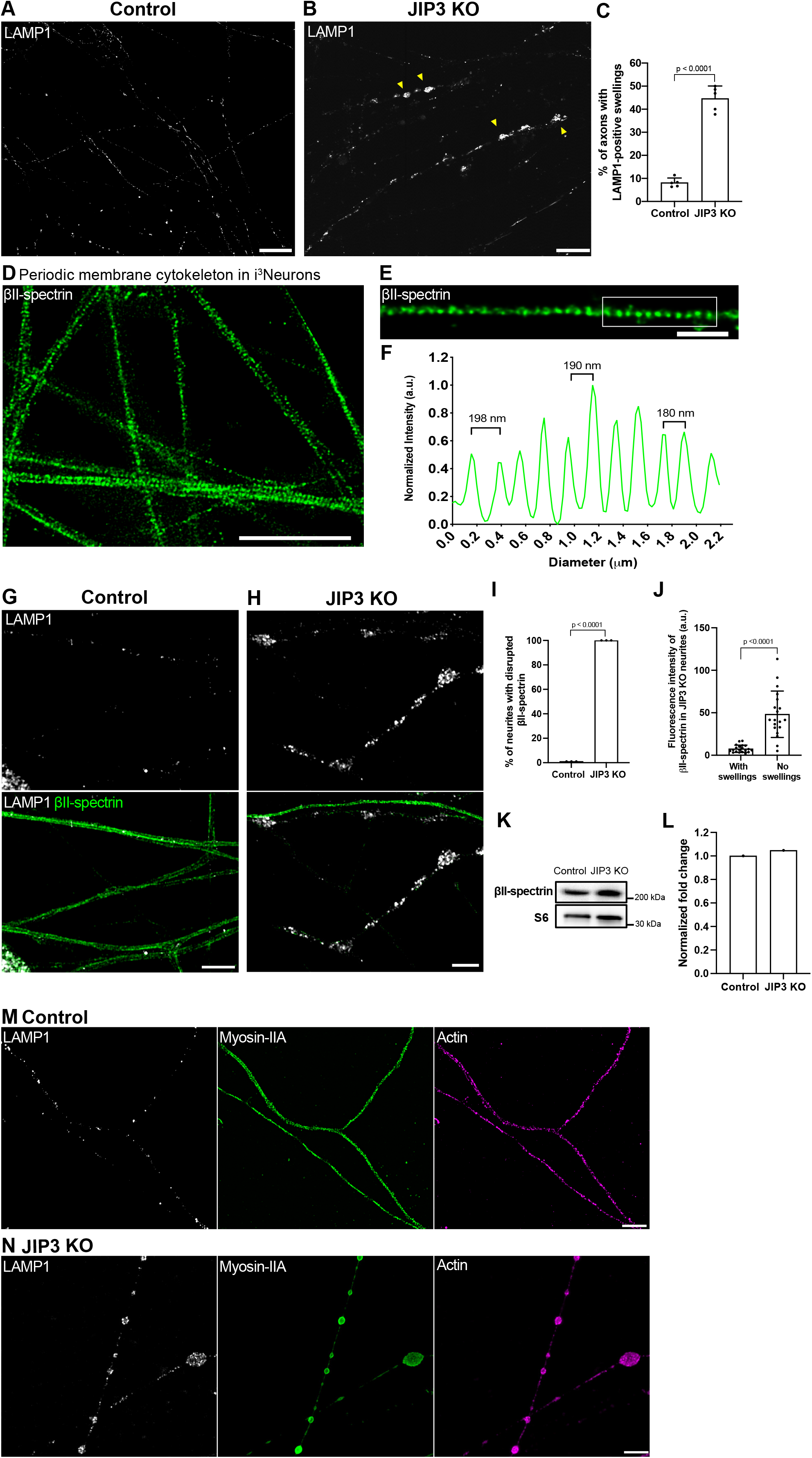
Lysosome-filled axonal swellings in JIP3 KO axons correlate with global disruption of the membrane periodic skeleton. (A and B) Airyscan imaging of control i^3^Neurons and JIP3 KO i^3^Neurons (13 days of differentiation). Yellow arrowheads highlight lysosome-positive axonal swellings in the KO neurons (scale bars, 15 μm). (C) Percentage of i^3^Neurons containing at least one lysosome-positive axonal swelling represented as mean ± SD, pooled from four independent experiments (n≥32 per experiment, 13 days of differentiation). (D and E) STED microscopy images of βII-spectrin immunofluorescence in the axons of control i^3^Neurons (day 17). (Scale bars, E: 5 μm; F: 1 μm). (F) Graph demonstrating the longitudinal distance between peaks in the βII-spectrin signal from the boxed region in (E). (G) Airyscan microscopy images of control i^3^Neurons show regular distribution of lysosomes (LAMP1, white) and intact periodic membrane skeletons (βII-spectrin, green). (H) JIP3 KO i^3^Neurons (day 15) exhibit disruption in the spectrin periodicity in axons positive for lysosome accumulations (scale bars, 5 μm). (I) Percentage of swollen axons with disrupted periodic membrane skeleton represented as mean ± SD pooled from three independent experiments (≥20 axons analyzed per experiment). Lysosomes and the periodic membrane skeleton were labeled with LAMP1 and βII-spectrin antibodies, respectively. (J) Graph depicting the mean βII-spectrin fluorescence intensity of JIP3 KO neurites with and without lysosome-filled axonal swellings (≥35µm in length) represented as mean ± SD, pooled from three independent experiments (n≥20 in total). Note the possible contribution of some dendrites to the “No Swellings” group. (K and L) Immunoblots showing levels of βII-spectrin in control and JIP3 KO i^3^Neurons (day 15); ribosomal protein S6 was used as loading control (K), and their normalized expression levels is shown in (L). (M) STED microscopy images of myosin-II filaments (green) show a periodic distribution similar to that of F-actin (magenta) in control i^3^Neurons. (N) Myosin-II and actin periodicity is lost in JIP3 KO i^3^Neurons (day 15) and both were enriched at the lysosome-positive axonal swellings (white). Lysosomes and myosin-II filaments were labeled with antibodies against LAMP1 and non-muscle myosin-IIA respectively, while rhodamine phalloidin was used to label F-actin. Scale bars, 5 μm. p-values were calculated using two-tailed Student’s t-test.

We next tested whether the axons of i^3^Neurons develop a membrane associated periodic skeleton similar to that observed in rodent neuron primary culture models (He et al., 2016; Xu et al., 2013). Stimulated Emission Depletion (STED) super-resolution fluorescence microscopy of control i^3^Neurons after labeling with antibodies against the C-terminus of βII-spectrin, revealed a periodic spectrin lattice (180-200nm intervals) (Figure 1D-F), in agreement with previous studies in other neuron culture systems (Xu et al., 2013; Zhong et al., 2014). In the control i^3^Neurons, this periodic spectrin organization was apparent by 9 days of differentiation and then persisted as the neurons aged (out to 21 days in this study; Figure 1D). This lattice was also observed in JIP3 KO neurons by 9 days of differentiation, a stage at which they have undergone extensive axon growth but do not yet display lysosome accumulations (Supplementary Figure 1A). In contrast, in older JIP3 KO i^3^Neurons that developed lysosome-filled axonal swellings, the axonal spectrin lattice was disrupted (Figure 1G-J). This striking disruption was not limited to the local site of lysosome accumulation but occurred throughout the entire affected axon (Figure 1G-J). This phenotype was even observed in axons with very sparse swellings (Supplementary Figure 1B). No difference was observed in the overall protein levels of βII-spectrin between control and JIP3 KO neurons (Figure 1K and L). The apparent loss of βII-spectrin signal in the microscopy experiments therefore most likely reflects the dispersal and washout of unassembled spectrin under the extraction conditions employed to enable super-resolution imaging of the rings formed by βII-spectrin. Consistent with the disruption of the spectrin periodic skeleton, the periodic organization of axonal F-actin was also lost and F-actin instead accumulated at the swellings (Figure 1M and N). Collectively, this data indicates that the axonal actin-spectrin lattice still formed in JIP3 KO neurons but later disassembled in parallel with the development of lysosome accumulations.

Non-muscle myosin-II associates with F-actin within the actin-spectrin lattice and controls axon radial contractility (Costa et al., 2020; Wang et al., 2020). We therefore examined the impact of the absence of JIP3 on the sub-cellular localization of myosin-II (Figure 1M and N). In control i^3^Neurons, myosin-II filaments displayed a periodic pattern with occasional gaps along the axonal shaft (Figure 1M). This periodicity was completely lost in the axons of JIP3 KO i^3^Neurons that had lysosome-filled axonal swellings (Figure 1N). In these axons, the myosin-II signal, like F-actin, was instead most prominent within the swellings.

### Lysosome-filled axonal swellings coincide with sites of abnormal microtubules organization

The assembly of the actin/spectrin-based axonal periodic scaffold is dependent on intact microtubules (Zhong et al., 2014). Given the drastic disruption of this scaffold in the swollen axons of JIP3 KO i^3^Neurons (as well as the essential role for microtubules in long range axonal transport of organelles), we next investigated microtubule organization. Remarkably, the sites at which lysosomes accumulate in JIP3 KO axons coincided with local disorganization of microtubules, as assessed by α-tubulin immunofluorescence (Figure 2A-C). The disorganization of microtubules was further exacerbated in older JIP3 KO i^3^Neurons (Figure 2D-F). The majority of microtubules within the swellings were bent and looped, while axonal regions immediately adjacent to the swellings contained microtubules that were organized in the typical parallel bundles of control axons (Figure 2B). This disruption of the normal parallel arrangement of microtubules throughout axons was even more striking in cultures of JIP3+JIP4 double KO i^3^Neurons, where, as previously reported (Gowrishankar et al., 2021), lysosome-filled swellings were larger and more abundant (Figure 2G-I). Of note, while lysosomes strongly accumulate in JIP3 KO i^3^Neurons and JIP3+JIP4 double KO i^3^Neurons, mitochondria and synaptic vesicles are also present in these lysosome-positive swellings, but to a much lesser degree (Supplementary Figure 2A-F).

**Figure 2:**
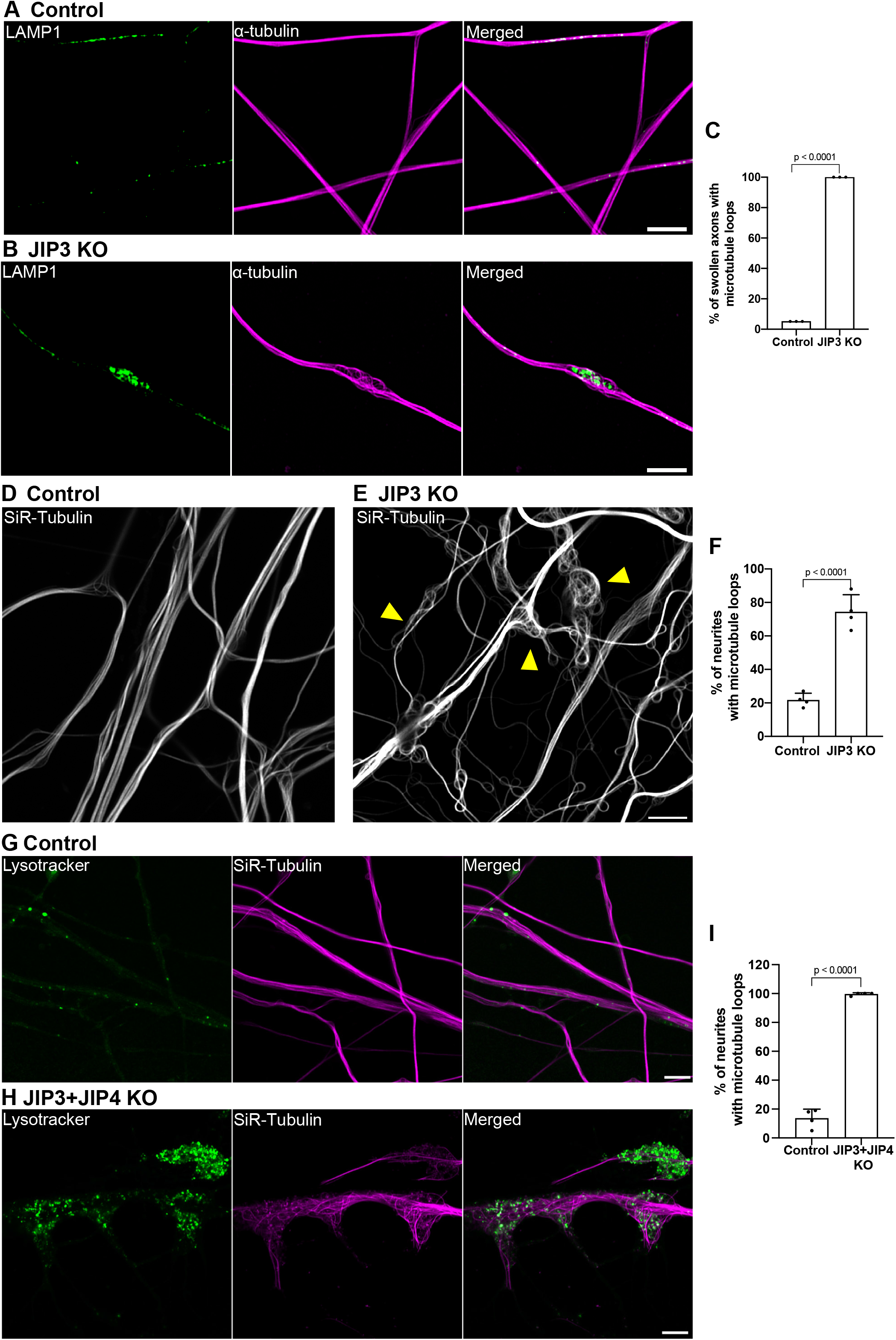
Lysosome-filled axonal swellings in JIP3 KO axons coincide with sites of abnormal microtubule organization. (A and B) Airyscan imaging of LAMP1 (green) and α-tubulin (magenta) immunofluorescence in control and JIP3 KO neurons (day 13) respectively (scale bars, 5 μm). (C) Percentage of swollen axons with abnormally looped microtubules presented (mean ± SD; pooled from three independent experiments, ≥30 axons per experiment). (D-E) Control and JIP3 KO neurons after 22 days of differentiation labeled for microtubules (SiR-tubulin; yellow arrowheads highlight examples of severe microtubule looping; scale bars, 5 μm). (F) The percentage of neurites (mean ± SD) with disorganized microtubules pooled from four independent experiments of 22 day old cultures (≥34 neurites per experiment). (G and H) Lysotracker (green) and SiR-Tubulin (magenta) in control and JIP3+JIP4 double KO i^3^Neurons (day 12) respectively (scale bars, 5 μm). (I) The percentage of neurites (mean ± SD) with disorganized microtubules pooled from four independent experiments (≥22 neurites per experiment). p-values in all experiments were calculated using two-tailed Student’s t-tests.

Microtubules undergo several distinct post-translational modifications (Baas et al., 1993; Baas and Black, 1990; Nirschl et al., 2017), which reflect microtubule age and the activities of multiple tubulin-modifying enzymes (Janke and Magiera, 2020; Park and Roll-Mecak, 2018). Immunofluorescent labeling revealed acetylation of axonal microtubules in both control and JIP3 KO i^3^Neurons, and there were no noticeable differences in the acetylation status of looped versus parallelly-organized microtubules in the KO neurons (Figure 3A-C). In contrast, when microtubule tyrosination status was examined, the looped microtubules were found to be predominantly detyrosinated (Figure 3D-F). While the mechanisms that underlie the relationship between looping and detyrosination of microtubules remain uncertain, it is possible that detyrosinated microtubules are more stable, potentially through inhibited binding of the depolymerizing kinesin (MCAK) to microtubules (Peris et al., 2009; Sirajuddin et al., 2014). In addition, as CAP-Gly domain containing proteins, such as the p150^Glued^ subunit of dynactin, prefer tyrosinated microtubules and tyrosinated microtubules have been proposed to be preferred tracks for initiating movement of axonal LAMP1-positive organelles (lysosomes) (Nirschl et al., 2016; Peris et al., 2006), accumulation of detyrosinated microtubules within axonal swellings could have an impact on lysosome transport at such sites.

**Figure 3:**
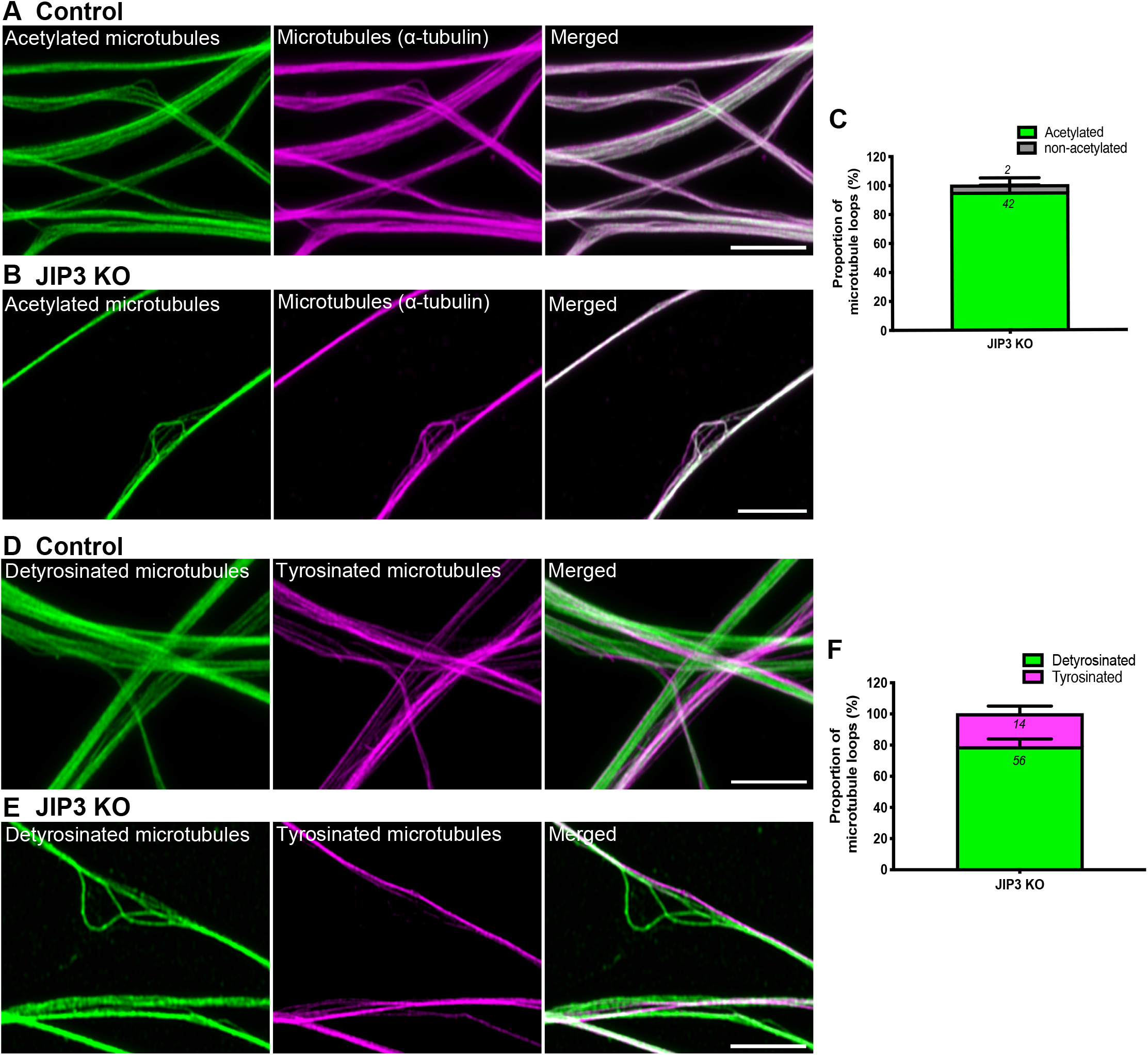
Microtubule loops in JIP3 KO axonal swellings are primarily detyrosinated. (A and B) Airyscan microscopy images of acetylated-α-tubulin, (green) and total microtubules (magenta) in control and JIP3 KO i^3^Neurons (day 13) respectively. Scale bars, 5 μm. (C) Fraction of acetylated and non-acetylated microtubules in JIP3 KO i^3^Neurons (pooled data from 2 experiment with ≥19 swellings analyzed per experiment). (D and E) Airyscan microscopy images of neurites from both control (D) and JIP3 KO i^3^Neurons (day 13) (E) consist of parallel microtubule bundles that are either detyrosinated (green) or tyrosinated (magenta) in control and JIP3 KO i^3^Neurons respectively. Scale bars, 5 μm. (F) Fraction of detyrosinated and tyrosinated microtubules in JIP3 KO i^3^Neurons (pooled from three independent experiments with ≥18 swellings analyzed per experiment).

### Axonal swellings are highly dynamic

We next performed long-term live cell imaging of lysosomes and microtubules in JIP3 KO i^3^Neurons to examine the dynamics of the axonal swellings. For the purpose of microtubule visualization, we employed low concentrations of the SiR-tubulin dye. This probe yielded a fluorescent microtubule pattern highly similar to the pattern of anti-α tubulin immunofluorescence (Figure 2A and B) including the appearance of microtubule loops within axonal swellings (Figure 2B, Figure 2D and E). Time-lapse imaging of JIP3 KO neurons stably expressing LAMP1-GFP (lysosome marker) and labeled with SiR-tubulin revealed that lysosome-filled axonal swellings form and resolve over a time scale of several hours (3.77 ± 1.71 hours; Figure 4A-D, Supplementary Movie S1). The close spatial and temporal relationship between lysosome accumulation and microtubule looping within these axonal swellings prevented definitive conclusions regarding a cause-effect relationship between these events (Figure 4B and C).

**Figure 4:**
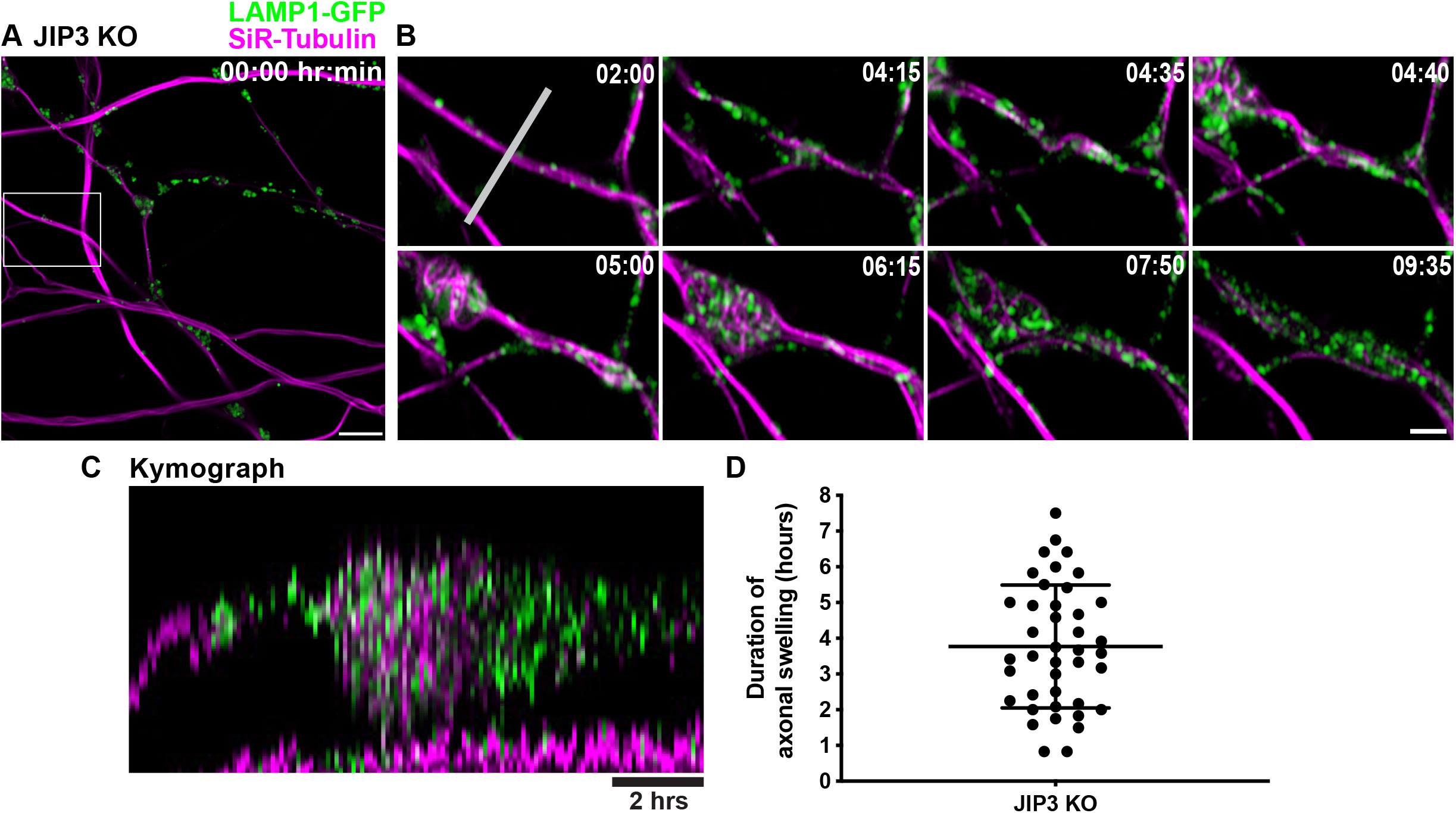
Lysosome-positive axonal swellings are dynamic in JIP3 KO neurons. (A) Representative image of JIP3 KO i^3^Neurons (day 15) stably expressing LAMP1-GFP (green) and labeled with SiR-tubulin (magenta) at the beginning of a 12-hour time lapse imaging time course (Airyscan microscopy). Scale bar, 5 μm. (B) Images from the boxed area in (A) at the indicated time points. Scale bar, 2 μm. (C) Kymograph of the region (marked by white line in B; scale bar, 2 hours). (D) Scatter dot plot (mean ± SEM) depicting the duration of 41 swellings pooled from 7 independent experiments.

To test whether microtubule perturbations can directly influence the focal accumulation of lysosomes, we acutely treated JIP3 KO i^3^Neurons with low doses of taxol (10nM or 20nM) for 1 hour, as higher concentration (1 µM) of taxol showed apparent toxicity in these neurons. We found that low doses of taxol efficiently mobilized large fractions of lysosomes from the axonal swellings within a period of 1 hour (Supplementary Figure 3A-B). These results further support a coordinative role for JIP3 in the maintenance of lysosome transport and microtubule stability.

To test the specificity of the phenotypes observed in JIP3 KO i^3^Neurons, we performed genome editing on JIP3 KO iPSCs to remove the 1 bp insertion in our JIP3 KO line and thus restore JIP3 expression (JIP3 Rescue). JIP3 expression was fully restored in the neurons derived from these JIP3 Rescue iPSCs (Figure 5A).

**Figure 5:**
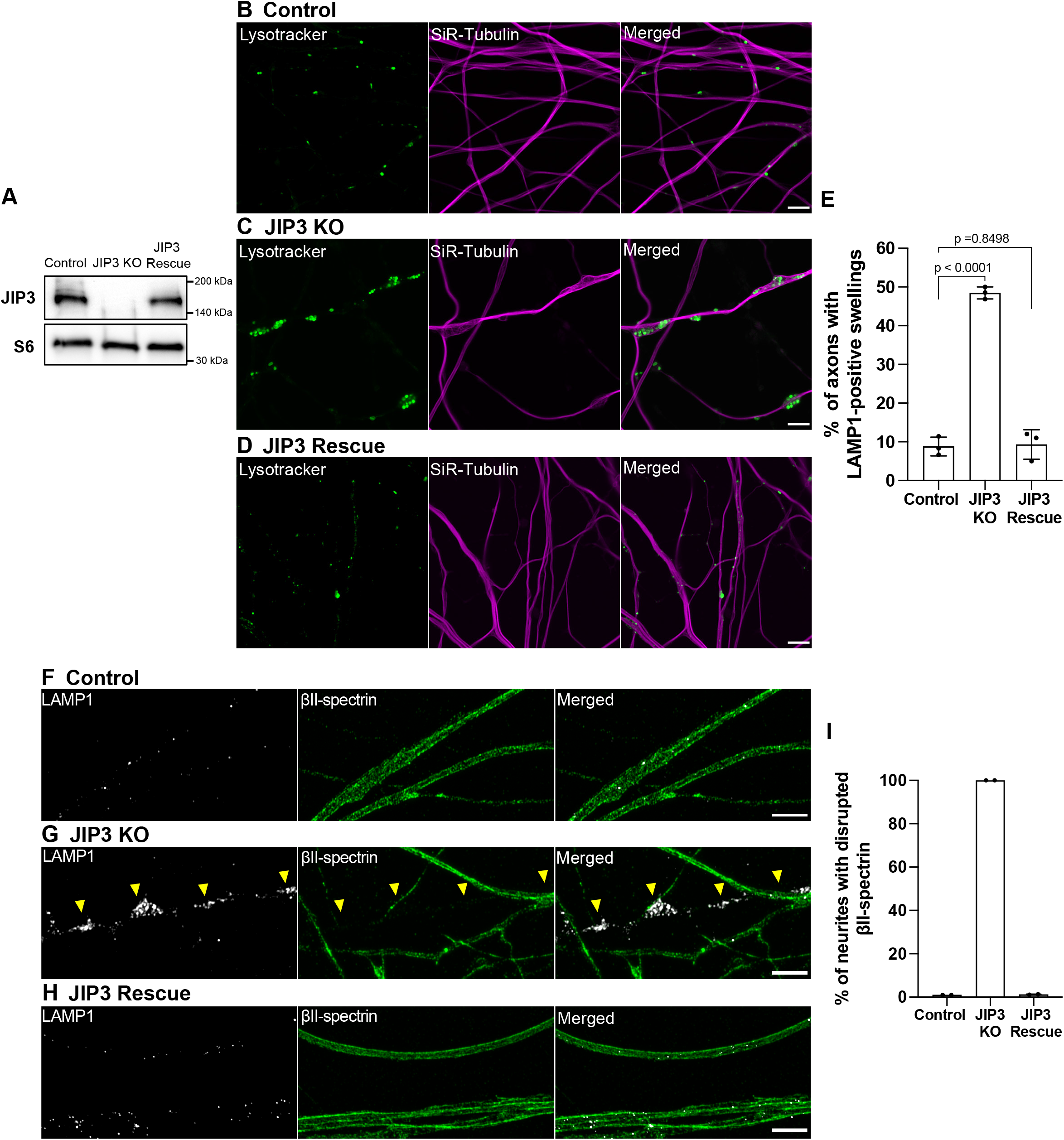
Rescue of JIP3 KO phenotypes. (A) Immunoblots showing levels of JIP3 in control, JIP3 KO and JIP3 Rescue (gene-edited to WT) i^3^Neurons (day 17). Ribosomal protein S6 immunoblots served as loading controls. (B-D) Airyscan imaging of Lysotracker (green) and Sir-tubulin (magenta) in control (B), JIP3 KO (C) and JIP3 Rescue (D) i^3^Neurons (day 17) respectively (scale bars, 5 μm). (E) Percentage of i^3^Neurons containing at least one lysosome-positive axonal swelling represented as mean ± SD, pooled from three independent experiments (n≥20 per experiment, 17 days of differentiation). (F-H) Airyscan microscopy images of lysosomes (white) and βII-spectrin (green) in control(F), JIP3 KO (G) and JIP3 Rescue (H) i^3^Neurons (Day 17), respectively. (scale bars, 5 μm). Lysosomes and the periodic membrane skeleton were labeled with LAMP1 and βII-spectrin antibodies, respectively. (I) Percentage of swollen axons with disrupted periodic membrane skeleton represented as mean ± SD pooled from two independent experiments (≥8 analyzed per experiment). p-values were calculated using two-tailed Student’s t-tests.

Furthermore, these JIP3 Rescue i^3^Neurons did not exhibit lysosome-positive focal swellings nor any disruptions to the microtubule organization (Figure 5B-E). In addition, the disruptions to the axonal spectrin lattice in swollen JIP3 KO axons (Figure 1G-I) were not found in JIP3 Rescue i^3^Neurons, as indicated by the βII-spectrin staining (Figure 5F-J). Collectively, these results demonstrate that both the lysosome accumulations and the cytoskeletal disorganization arose specifically due to the loss of JIP3 rather than any off-target effect of the genome editing strategy that was employed to generate the JIP3 KO line.

## Discussion

Our investigation of the relationships between lysosome axonal transport and multiple components of the axonal cytoskeleton in neurons that lack JIP3, or both JIP3 and its paralogue JIP4, revealed that focal lysosome accumulations are accompanied by major disruptions in organization of the axonal membrane associated periodic skeleton. Furthermore, the accumulation of lysosomes in focal swellings of mutant neurons coincided both spatially and temporally with local microtubule disorganization. These observations reveal major unexpected relationships between axonal lysosome transport and the organization of multiple aspects of the axonal cytoskeleton. They furthermore raise new questions about the chain of events that link lysosome transport, microtubule organization and the cytoskeletal machinery that controls axon diameter.

Although impacts of JIP3 depletion on axonal lysosome abundance have been observed in multiple species (Drerup and Nechiporuk, 2013; Edwards et al., 2013; Gowrishankar et al., 2017), broader effects of JIP3 mutation on the axonal abundance of multiple organelles have also been reported (Brown et al., 2009; Byrd et al., 2001; Noma et al., 2017; Sato et al., 2015; Sure et al., 2018). Our new observations of the dramatic cytoskeleton disruption that correlates with focal lysosome accumulations in both JIP3 KO (and JIP3+JIP4 double KO) neurons provide a potential explanation for the broad requirement for JIP3/4 for the axonal transport of multiple organelles that would not require a direct involvement of JIP3/4 in the transport of each class of organelles.

The focal accumulation of lysosomes coincides spatially and temporally with local disorganization of microtubules in JIP3 KO i^3^Neurons and the overall severity of this phenotype is further exacerbated in the JIP3+JIP4 double KO neurons. Our initial interpretation of these lysosome phenotypes was that lysosomes fall off their microtubule tracks due to a loss of JIP3+4-dependent connections to motors. The surprising discovery that these sites of lysosome accumulation were accompanied both spatially and temporally with extensive looping of microtubules suggests a more complex relationship between lysosome transport and microtubule organization.

It was particularly striking that the dynamic formation of local axonal swellings with lysosome accumulation and microtubule looping in JIP3 KO neurons was accompanied by a widespread disruption of the integrity of the axonal membrane-associated periodic skeleton. This all-or-none effect is consistent with two distinct interpretations. One is that the focal swellings elicit changes that are propagated along the entire axonal shaft. Another is that loss of JIP3 results in an age-dependent disruption of the periodic actin/spectrin based scaffold that facilitates formation of the focal swellings where lysosomes accumulate. Interestingly, the assembly of the axonal membrane associated periodic skeleton is dependent on microtubule integrity (Qu et al., 2017; Zhong et al., 2014), suggesting that the cytoskeletal changes observed in JIP3+JIP4 KO neurons may be interconnected. Interaction between the periodic membrane skeleton and microtubules may occur via several linker proteins, including ankyrins, which bind both spectrin and microtubules (Bennett and Davis, 1981; Leterrier et al., 2011; Zhong et al., 2014). More recently, dynein has been implicated in the maintenance of the actin-spectrin cytoskeleton at the axon initial segment through endocytosis-related mechanisms(Torii et al., 2020). This provides another means through which the integrity of the actin-spectrin rings can be regulated by motor proteins. The elucidation of the precise sequence of events will require further experimentation.

The phenotypes that we observed in JIP3 KO axons are reminiscent of axonal beading that arises in response to multiple forms of axonal perturbation (Bar-Ziv et al., 1999; Fernandez and Pullarkat, 2010; Kilinc et al., 2009). It was recently shown that changes in the local tension of the axonal membrane leads to the propagation of such “beads” (Datar et al., 2019). Myosin-II filaments within the periodic membrane skeleton contribute to contractility and may control tension homeostasis along the axonal shaft (Wang et al., 2020). Hence, if the dilations are upstream events, the all-or-none disruption of the axonal periodic scaffold within swollen JIP3 KO axons may be a consequence of tensional instability along the axons.

Loss of JIP3 could also have signaling consequences that could propagate beyond its direct subcellular site of action. In addition to interactions with motors, JIP3 (and JIP4) also intersect with signaling in the JNK pathway by acting as a scaffold that regulates the subcellular position and activity of DLK and JNK (Drerup and Nechiporuk, 2013; Ghosh et al., 2011; Kelkar et al., 2000; Kulkarni et al., 2019). Additionally, transport defects could impede the ability of lysosomes to act as sites for nutrient and growth factor-dependent activation of the mTORC1 signaling pathway (Ferguson, 2015; Liu and Sabatini, 2020).

In conclusion, our findings have revealed new reciprocal interrelations between lysosome transport and the structure and the dynamics of the axonal cytoskeleton. The stalling of lysosomes during their retrograde journey correlated with profound changes in axons. A priority for future work will be to determine cause-effect relationships between the multiple tightly linked changes that we have observed here. In addition to advancing our knowledge about fundamental aspects of cell function, these studies may provide new insight into mechanisms relevant to Alzheimer’s disease pathology as well as to human intellectual disabilities arising from mutations in the *MAPK8IP3/JIP3* gene (Iwasawa et al., 2019; Platzer et al., 2018).

## Materials and Method

### Human iPSC culture and neuronal differentiation

Human iPSCs were differentiated into cortical i^3^Neurons according to a previously described protocol based on the doxycycline inducible expression of Ngn2 (Fernandopulle et al., 2018). Briefly, the iPSCs were cultured on human embryonic stem cell (hESC)-qualified Matrigel (Corning) and fed with fresh mTeSR™1 medium (STEMCELL Technologies) on alternate days. Rho-kinase (ROCK) inhibitor Y-27632 (EMD Millipore, 10 μM) was added to the iPSC cultures on the first day of plating and replaced with fresh media without ROCK inhibitor on the following day. For neuronal differentiation, iPSCs were dissociated with Accutase (STEMCELL Technologies) and re-plated at a density between 1.5-3×10^5^ cells on matrigel-coated dishes in induction medium (KnockOut DMEM/F-12 (Thermo Fisher Scientific) containing 1% N2-supplement [Gibco], 1% NEAA [Gibco], 1% GlutaMAX [Gibco]) and 2 μg/mL doxycycline [Sigma]). After 3 days, pre-differentiated i^3^Neurons were dispersed using Accutase and plated on poly-L-ornithine (Sigma, 1 μg/ml) and laminin (Thermo Fisher Scientific, 10 μg/ml) coated 35 mm glass-bottom dishes (MatTek) or 6-well plates (Corning) for imaging and immunoblotting, respectively. These i^3^Neurons were cultured and maintained in cortical medium (induction medium supplemented with 2% B27 (Gibco), 10 ng/mL BDNF (PeproTech), 10 ng/mL NT-3 (PeproTech) and 1 μg/mL laminin). Fresh cortical media was added to the existing media every 5 days. The iPSCs and i^3^Neurons were kept at 37°C with 5% CO_2_ in an enclosed incubator.

### CRISPR-Cas9 mediated rescue of JIP3 KO iPSCs

A CRISPR-based homologous recombination strategy was used to correct the 1 bp insertion in the JIP3 KO iPSC line. Briefly, 1x 10^5^ JIP3 KO iPSCs were plated on Matrigel-coated 6-well plate and transfected the following day using the Lipofectamine Stem transfection reagent (Thermo Fisher Scientific) and the following two components: 3µg of px458 plasmid (Addgene plasmid #48138) containing a small guide RNA encoded within the following sense (5’CACCGGGCGGCGTGGTGGTGTTACC3’) and antisense (5’AAAC GGTAACACCACCACGCCGCCC 3’) sequences that was designed to selectively target the JIP3 KO sequence containing the 1 bp insertion and a 140 bp single stranded DNA oligonucleotide repair template (5µl of 100µM stock) that overlapped the gRNA-targeted region. The sequence of the DNA template is the following: 5’GCCGCGCTGGCGGCGGCGGTGGCCGCGATGATGGAGATCCAGATGGACGAGGGCGGCGGCGTGGTGGTGTACCAGGACGACTACTGCTCCGGCTCGGTGATGTCGGAGCGGGTGTCGGGCCTGGCGGGCTCCATCTACCG 3’. Transfected (GFP-positive) cells were enriched by fluorescence activated cell sorting (FACS) 2 days later. Sorted cells were expanded and then serially diluted to yield single cell-derived clonal populations. 40 clones were selected and screened using PCR amplification of genomic DNA flanking the sgRNA target site followed by sequencing of the amplicons using the primers described in Gowrishankar et al. (2020).

### Live cell imaging and drug treatments

Live imaging of control, JIP3 KO and JIP3+4 KO i^3^Neurons (Gowrishankar et al., 2020) was performed on day 10-22 post-differentiation in cortical medium supplied with 5% CO_2_ and maintained at 37°C. I^3^Neurons stably-expressing LAMP1-GFP (Gowrishankar et al., 2020) or LysoTracker-labelled i^3^Neurons were used to visualize lysosome dynamics. For lysotracker and mitotracker labeling, i^3^Neurons were stained with 30 nM LysoTracker™ Green DND-26, 10 nM LysoTracker™ Deep Red or 100nM MitoTracker™ Deep Red (Thermo Fisher Scientific) for 3 mins, washed twice with fresh cortical media, and then imaged immediately. To label synaptic vesicles, i^3^Neurons were transfected with 4 μl Lipofectamine Stem Transfection Reagent (Thermo Fisher Scientific) in 200 μL OptiMEM medium mixed with 3 μg synaptophysin-GFP plasmid and incubated for 10 minutes. The Lipofectamine-DNA complex was added to the imaging dish, followed by media replacement with fresh cortical media on the following day. For Taxol treatment, i^3^Neurons were labeled with lysotracker as described above and then treated with 10nM or 1μM taxol (Sigma) for 1 hour.

### Immunofluorescence

For i^3^Neuron samples involving actin and spectrin staining, cells were fixed and extracted for 1 minute using a solution of 0.3% (v/v) glutaraldehyde and 0.25% (v/v) Triton X-100 in cytoskeleton buffer (CB, 10 mM MES pH 6.1, 150 mM NaCl, 5 mM EGTA, 5 mM glucose and 5 mM MgCl_2_), post-fixed for 15 min in 2% (v/v) glutaraldehyde in CB at 37°C, and then washed twice in phosphate-buffered saline (PBS) according to previously described protocol (Xu et al., 2013). For microtubule staining, cells were fixed and extracted for 15 minutes using a solution of 4% (v/v) paraformaldehyde, 0.2% (v/v) glutaraldehyde and 0.25% (v/v) Triton X-100 in CB at 37°C, and then washed twice in PBS. For removal of free aldehyde groups, cells were quenched with fresh 1 mg/ml sodium borohydride in CB (Sigma) for 10 mins, and then washed thrice in PBS. Cells were further blocked for 30 minutes in 5% bovine serum albumin (BSA, Sigma) in phosphate-buffered saline (PBS) and then incubated overnight at 4°C with the following appropriate antibodies: anti-JIP3 (Novus Biologicals, catalogue no. NBP1-00895, dilution 1:500); anti-α-tubulin (Sigma, catalogue no. T6199, dilution 1:500); anti-LAMP1 (Cell Signaling Technology, catalogue no. 9091, dilution 1:500) or (Developmental Studies Hybridoma Bank, clone 1D4B, dilution 1:500); anti-βII-spectrin (BD Transduction Laboratories, Clone 42/B-Spectrin II, dilution 1:250); anti-non muscle heavy chain of myosin-IIA (Sigma, catalogue no. M8064, dilution 1:500); anti-acetyl α-tubulin (Cell Signaling Technology, catalogue no. 5335, dilution 1:250); anti-detyrosinated α-tubulin (Abcam, catalogue no. ab48389, dilution 1:250); anti-tyrosinated α-tubulin (Millipore Sigma, clone YL1/2, dilution 1:250). Cells were washed with PBS thrice and incubated with Alexa Fluor-conjugated secondary antibodies (Thermo Fisher Scientific) for 1 hour at room temperature, followed by three washes in PBS. F-actin was visualized by Alexa Fluor 488 or rhodamine-conjugated phalloidin (Thermo Fisher Scientific, dilution 1:100).

### Immunoblotting

Control, JIP3 KO and JIP3+4 KO i^3^Neurons were grown on six-well plates (3 × 10^5^ cells/well). After post-differentiation in cortical media, i^3^Neurons (typically 17 days post-differentiation) were washed with ice-cold PBS and then lysed in lysis buffer (50 mM Tris pH 7.4, 150 mM NaCl, 1 mM EDTA, and 1% Triton X-100) supplemented with cOmplete™ EDTA-free protease inhibitor cocktail (Roche) and PhosSTOP phosphatase inhibitor cocktail (Roche), followed by centrifugation at 13,000xg for 6 min. The supernatant was collected and incubated at 95°C for 5 min in SDS sample buffer containing 1% 2-mercaptoethanol (Sigma). The extracted proteins were separated by SDS-PAGE in Mini-PROTEAN TGX precast polyacrylamide gels (Bio-Rad) and transferred to nitrocellulose membranes (Bio-Rad) at 100V for 1 hour or 75V for 2 hours (for high molecular weight proteins: >150kDa). Subsequently, the nitrocellulose membranes were blocked for 1 hour with 5% non-fat milk (AmericanBIO) in TBST (tris-buffered saline [TBS] + 0.1% tween 20), then incubated overnight at 4°C with primary antibodies: anti-S6 Ribosomal Protein (S6, Cell Signaling Technology, catalogue no. 2217, dilution 1:2500); anti-βII-spectrin (BD Transduction Laboratories, Clone 42/B-Spectrin II, dilution 1:1000).

Subsequently, the nitrocellulose membranes were washed 3 times (10 minutes each) in TBST and probed by incubation for 1 hour with the secondary antibodies conjugated with horseradish peroxidase. The membranes were then washed three times (15 minutes at room temperature each), developed using Pierce™ ECL western blotting substratum (Thermo Fisher Scientific) and imaged by a Versa-Doc imaging system (Bio-Rad).

### Fluorescence microscopy

Two types of high-resolution microscopes were used in this study. The LSM 880 inverted confocal laser scanning microscope with Airyscan (Carl Zeiss Microscopy) accompanied with 63×/1.40 numerical aperture (NA) plan-apochromat differential interference contrast (DIC) oil immersion objective and 32-channel gallium arsenide phosphide (GaAsP)-photomultiplier tubes (PMT) area detector and 488 nm, 561 nm and 633 laser lines was used in this study. Images were acquired and processed using ZEN imaging software (Zeiss). The Leica TCS SP8 gated STED super-resolution confocal microscope (Leica Microsystems) is coupled with Leica harmonic compound (HC) plan apochromatic (PL APO) 100×/1.40 oil STED objective and Leica gated HyD hybrid detector. Briefly, a white light excitation laser accompanied with 592 nm, 660 nm and 775 nm depletion lasers was used in this study. Images were acquired using LAS X software (Leica Microsystems) and final images were deconvolved using Huygens deconvolution software (Huygens Essentials, Scientific Volume Imaging).

### Quantification and statistical analysis

Images were pseudocolor-coded, adjusted for brightness and contrast, cropped and/or rotated using the open-source image processing software FIJI (ImageJ) (Schindelin et al., 2012). Dendrites and axons were identified by visual tracking of the length of the neurite. Percentage of neurites were quantified using the FIJI plugin “NeuronJ” and/or the FIJI segmented lines + ROI manager tool to determine the total number of neurites. For quantification of βII-spectrin intensity, neurites of at least 35 μm in length were semi-automatically traced and quantified using “NeuronJ” to determine the mean fluorescence value (8-bit) of the traced segments. In the case of JIP3 KO i^3^Neurons where βII-spectrin intensity is markedly reduced, LAMP1 staining was used to trace the swollen axons as described above. The traced segments were then overlayed to the βII-spectrin channel for subsequent quantification.

Western blot data were processed using Image Lab software (Bio-Rad) and quantified using the “Gels” ImageJ plugin. The methods for statistical analysis and sizes of the samples (n) are specified in the results section or figure legends for all of the quantitative data. Student’s t test or Mann-Whitney test was used when comparing two data sets. Differences were accepted as significant for P < 0.05. Prism version 8 (GraphPad Software) was used to plot, analyze and represent the data.

## Acknowledgements

We are grateful to Michael Ward (NINDS) for his contribution of the human iPSCs with doxycycline-inducible Ngn2 expression and his generous advice on their use. Agnes Roczniak-Ferguson played a key role in establishing iPSC genome editing and neuronal differentiation in our lab. Christopher Lovejoy provided helpful feedback on analysis of cytoskeleton phenotypes. This research was supported in part by the Dementia Discovery Foundation (SMF and PDC), the NIH (NS36251 to PDC; AG062210 TO SMF), and the Kavli Foundation (PDC). The authors do not have any conflicts to declare.

## Author contributions

NBMR, LLL, PDC and SMF designed experiments. NBMR and LLL performed all experiments. SG developed key reagents. NBMR, PDC and SMF prepared the manuscript.

## Figure Legends

**Supplementary Figure 1: βII-spectrin organization in JIP3 KO i**^3^**Neurons**

(A) Airyscan microscopy images of young JIP3 KO i^3^Neurons (day 9) with no lysosome-positive swellings (LAMP1, white) display intact periodic membrane skeleton (βII-spectrin, green). (B) The global disruption of the periodic membrane skeleton shown in Figure 1H is also seen in JIP3 KO i^3^Neuron (day 13) with very sparse lysosome-positive axonal swellings. Lysosomes and the periodic membrane skeleton were labeled with LAMP1 and βII-spectrin antibodies, respectively. Scale bars, 5 μm.

**Supplementary Figure 2: In addition to lysosomes, synaptic vesicles and mitochondria are also present within axonal swellings in the JIP3 KO and JIP3+JIP4 KO i**^3^**Neurons**

(A and B) Representative Airyscan microscopy images of lysosomes (green) and mitochondria (magenta) in control (A) and JIP3 KO (B) i^3^Neurons (day 15). Scale bars, 5 μm. Note that while lysosomes strongly accumulate in JIP3 KO i^3^Neurons, mitochondria are also present in these lysosome-positive swellings, but to a much lesser degree. (C, D) Airyscan microscopy images of synaptic vesicles (green) and lysosomes (magenta, lysotracker) control and JIP3+JIP4 KO i^3^Neurons respectively (10 days of differentiation). (E,F) Mitochondria localization in control versus JIP3+JIP4 KO i^3^Neurons (Airyscan images, day 10). Scale bars, 10 μm.

**Supplementary Figure 3: Effect of taxol on axonal swelling in JIP3 KO i**^3^**Neurons**

(A) Representative Airyscan microscopy image of JIP3 KO i^3^Neuron with lysosome-positive axonal swellings (lysotracker, white). Scale bar, 5 μm. (A’) Images from the boxed area in (A) at the indicated time points show dispersion of lysosomes from focal swellings (yellow arrowheads) when treated with low doses of taxol (10nM). Scale bar, 2 μm. (B) Percentage of neurites (mean ± SD) showing dispersion of lysosomes from focal swellings in JIP3 KO i^3^Neurons in the indicated conditions (pooled from at least three independent experiments with ≥3 swellings analyzed per experiment).

**Supplementary Movie 1: Dynamic formation and resolution of axonal swellings in JIP3 KO i**^3^**Neurons**

JIP3 KO i^3^Neurons (day 15) stably expressing LAMP1-GFP were labeled with SiR tubulin to mark lysosomes and microtubules, respectively. Images were acquired at 5-minute intervals over a period of 12 hours using Airyscan microscopy. Scale bar, 5 µm. Display rate is 5 frames/sec.

