## Supplementary figures and images for "JIP3 links lysosome transport to regulation of multiple components of the axonal cytoskeleton"

### Supplemental Data

#### Supplementary Figure 1

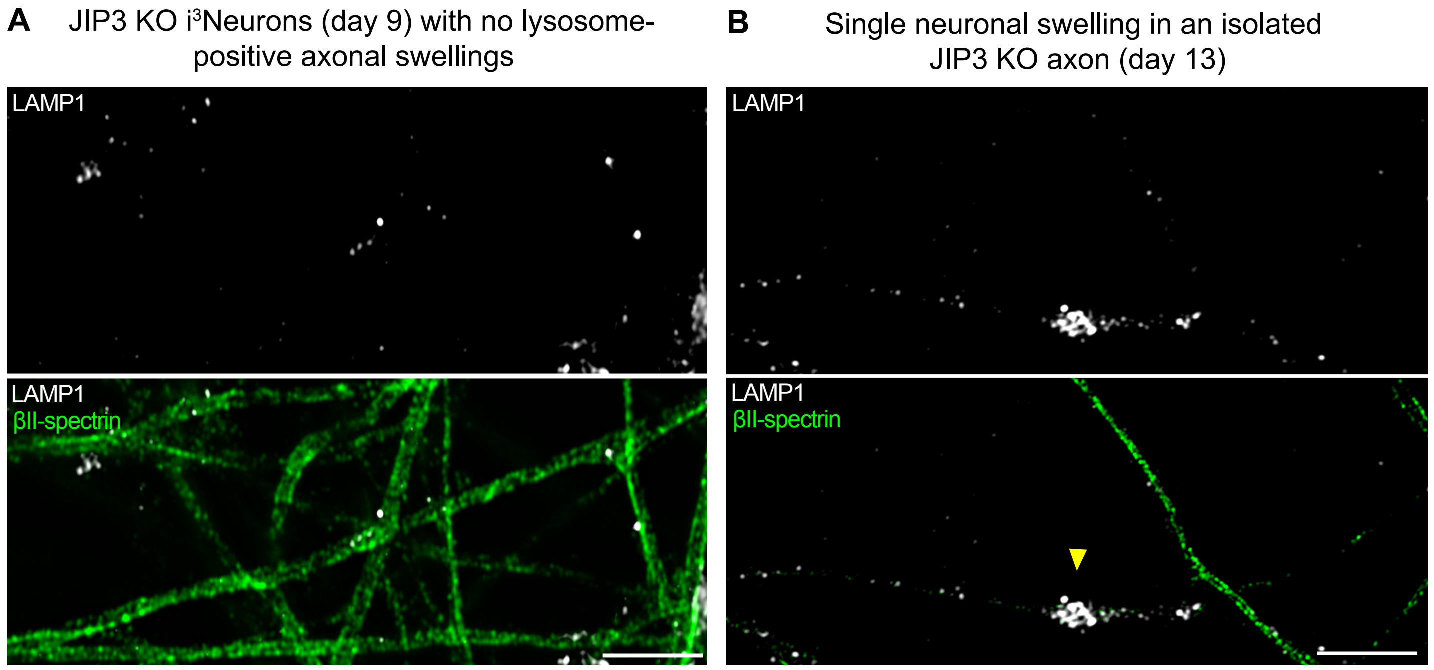

#### Supplementary Figure 2

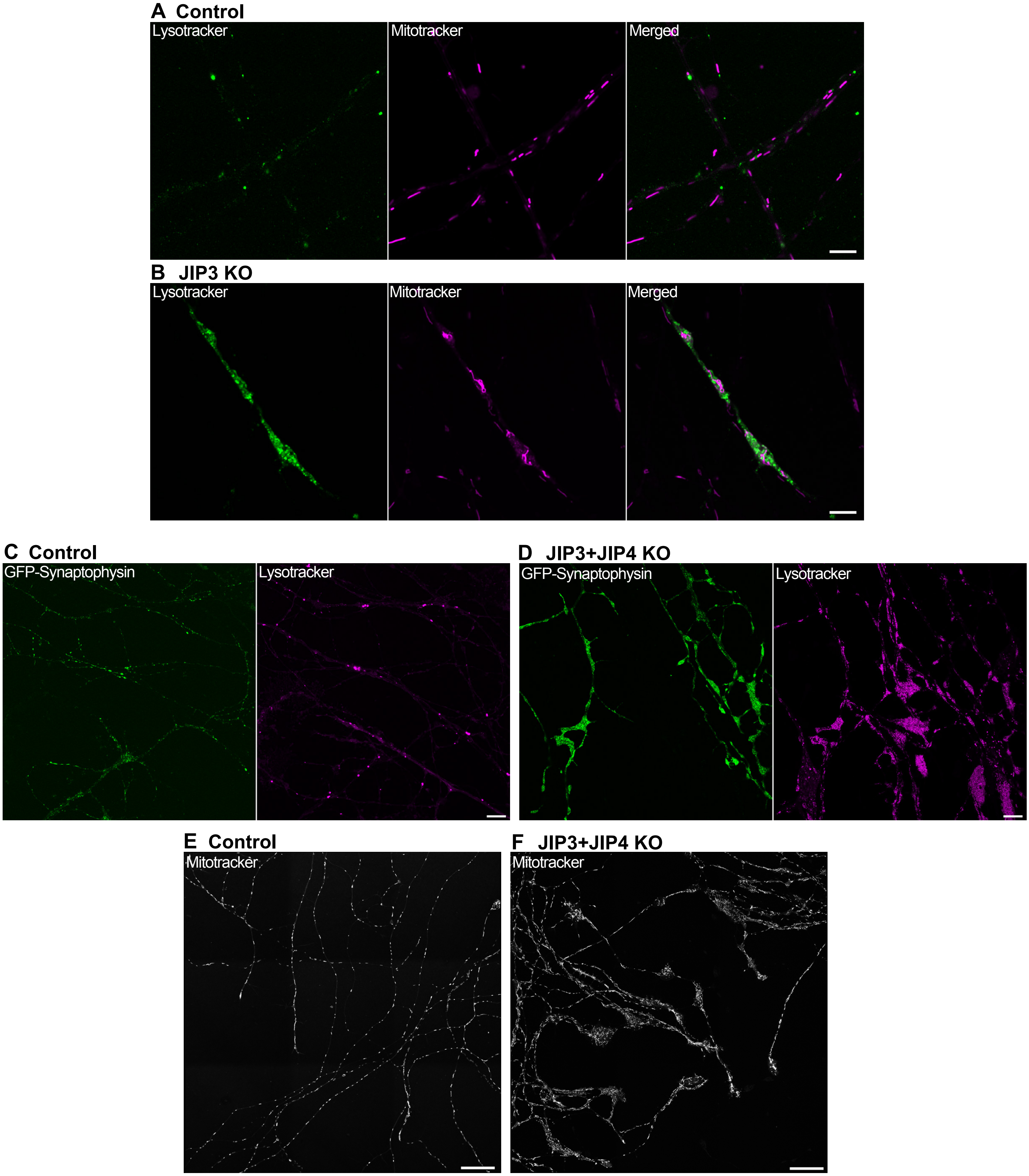

#### Supplementary Figure 3

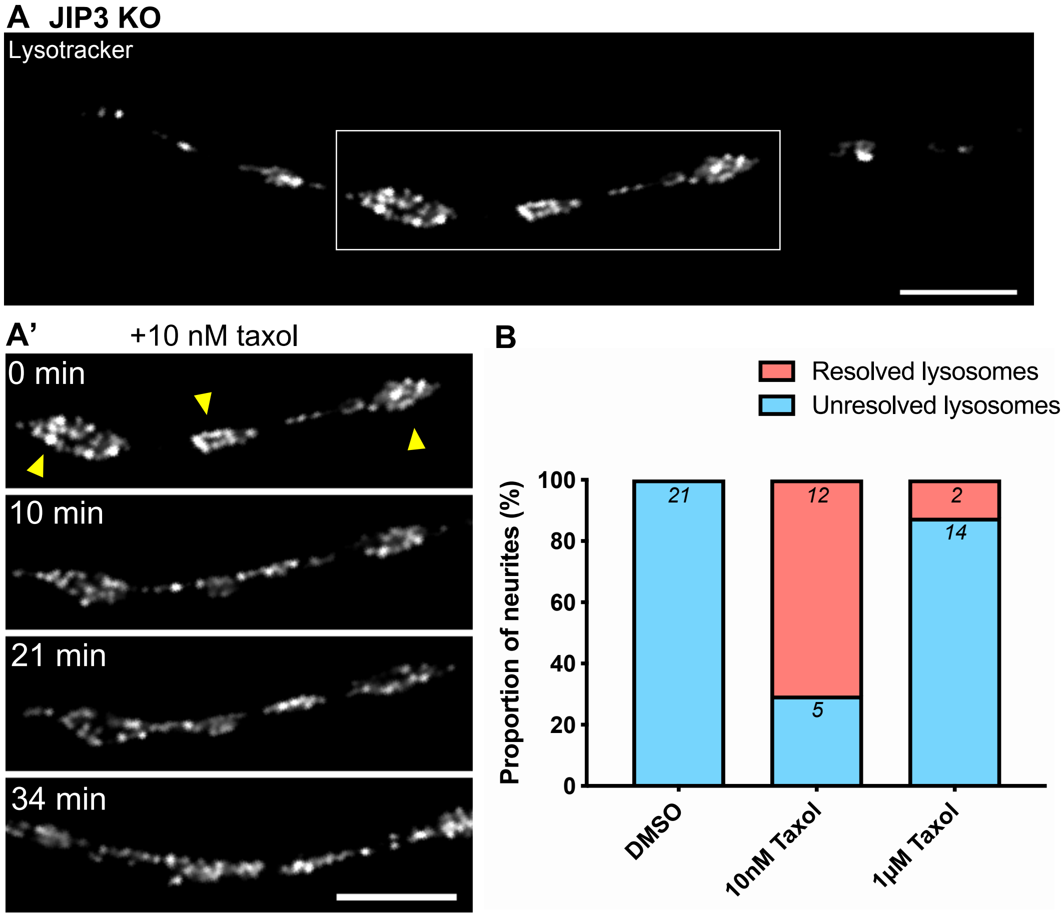
